# Time-scales modulate optimal lysis-lysogeny decision switches and near-term phage fitness

**DOI:** 10.1101/2021.06.21.449334

**Authors:** Shashwat Shivam, Guanlin Li, Adriana Lucia-Sanz, Joshua S. Weitz

## Abstract

Temperate phage can initiate lysis or lysogeny after infecting a bacterial host. The genetic switch between lysis and lysogeny is mediated by phage regulatory genes as well as host and environmental factors. Recently, a new class of decision switches was identified in phage of the SPbeta group, mediated by the extracellular release of small, phage-encoded peptides termed arbitrium. Arbitrium peptides can be taken up by bacteria prior to infection, modulating the decision switch in the event of a subsequent phage infection. Increasing concentration of arbitrium increases the chance that a phage infection will lead to lysogeny, rather than lysis. Although prior work has centered on the molecular mechanisms of arbitrium-induced switching, here we focus on how selective pressures impact the benefits of plasticity in switching responses. In this work, we examine the possible advantages of near-term adaptation of communication-based decision switches used by the SPbeta-like group. We combine a nonlinear population model with a control theoretic approach to evaluate the relationship between a putative phage reaction norm (i.e., the probability of lysogeny as a function of arbitrium) and the near-term time horizon. We show the adaptive potential of communication-based lysis-lysogeny responses and find that optimal switching between lysis to lysogeny increases near-term fitness compared to fixed responses. We further find that plastic responses are robust to the inclusion of cellular-level stochasticity. These findings provide a principled basis to explore the long-term evolution of phage-encoded decision systems mediated by extracellular decision-signaling molecules, and the feedback between phage reaction norms and ecological context.

## 1 Introduction

Upon infection, temperate phages like P1, P2 [6], phage *λ* [15] and *µ* [14] can initiate a lytic or lysogenic pathway whereas virulent phages can only initiate the lytic pathway. In a lytic pathway, the virus hijacks host cellular machinery to produce viral RNA and proteins, replicate the viral genome, assemble mature virus particles, lyse (and kill) the host cell, and ultimately, release virus particles [27] into the extracellular environment. In a lysogenic pathway the viral genome is integrated into the host genome. The stably integrated viral genome, i.e., the ‘prophage’, is then replicated along with the host [17] where the prophage is vertically transmitted to progeny cells. The integration of the phage genome often confers benefits to the lysogen, including immunity to infection and direct benefits to growth [1, 7, 13, 9]. Finally, the prophage can be induced, leading to the excision of the viral genome, and the re-initiation of the lytic pathway, including the release of virus particles [20].

Despite the existence of multiple pathways for propagation of viral genomes, the evolutionary ‘success’ of a phage has often been measured in terms of the quantity of viral particles generated via lysis or in terms of phage abundance in absolute terms [8]. However, contributions of viral replication to fitness need not include production of virus particles, at least not in the short term. As such, here we use a cell-centric metric for quantifying and characterizing viral fitness [25]. In doing so, we define the near-term Malthusian fitness as the growth rate of phage-infected cells, irrespective of whether these cells are part of the lysogenic or lytic pathways. This fitness depends not only on phage strategies but on ecological context.

In prior work using this cell-centric metric [5, 23, 24, 25, 16], host abundances were shown to influence the invasibility of phage. Increasing susceptible host densities increases the chance that phage particles released via lysis encounter and infect new hosts. In contrast, given low susceptible densities, phage that integrate into hosts can proliferate as lysogens while competing with fewer cells. These findings, framed in terms of the invasion fitness criteria for temperate phage, suggest that lysogeny can be an adaptive benefit in the near-term, particularly when host cell densities are low. Extrapolating from these prior findings, an intracellular sensing mechanism that allows phage to adjust life cycle decisions after infection given variation in host cellular density could be of adaptive benefit.

The study of the switch between lysis and lysogeny remains an important field of study given the prevalence of prophages in bacterial chromosomes and their impact on bacterial behavior, pathogenicity and evolution [17, 1, 7, 13, 9, 21]. Seminal work showed that the population-level frequency of lysogeny in phage *λ* increased with increasing ratios of phage to hosts [15]. Hence, when phage were relatively more abundant than bacterial hosts, phage infections tended to lead to lysogeny, rather than lysis. Recent work using single-cell imaging methods revealed that the probability of lysogeny increases with increasing cellular multiplicity of infection (from ~20% given a single phage to ~80% given five coinfecting phage) [28, 12]. These shifts in cellular fate reflect a combination of interactions between phage-specific gene regulatory circuits and the ecological context (which drives the multiplicity of infection).

It is increasingly apparent that lysis-lysogeny decisions are also mediated by small signaling molecules released to the environment. For example, the phage *VP882* uses the quorum sensing molecule acyl-homoserine lactone (Als) produced by the host, *Vibrio cholerae* to activate genes responsible for lysis [22]. In this case, an increase in host density translates to an increase in Als levels in the medium, and the probability of lysogeny decreases. Likewise, phage of the SPbeta group utilize the quorum sensing molecule arbitrium for lysis-lysogeny decisions [11]. Upon infection, the phage genetic arbitrium system regulates the release of a small peptide from infected cells that increase the probability that future infections are lysogenic, rather than lytic. In the absence of arbitrium in the environment, a receptor activates *aimX* expression, which in turn directs the phage to a lytic cycle. Instead, in the presence of arbitrium, the receptor binds to the peptide and the expression of *aimX* is repressed, which preferentially initiates the lysogenic cycle. A recent modelling study [10] deduced that the communicating phage outcompete non-communicating phage, particularly when host densities become low. Selection of lysis when susceptible host cells are abundant and lysogeny when susceptible cells become scarce is hypothesized to increase the long-term growth rate of phage that utilize a communicating strategy. Arbitrium systems may also include additional quorum sensing circuits that increase the efficiency of switching between lysis and lysogenic pathways [4]. As a final example, the phage phi3T employs two communication systems, arbitrium for lysis-lysogeny and Rap*φ*-Phr*φ* that downregulates host defenses and confers a fitness advantage to lysogenized cells. These examples demonstrate a connection between the cellular fate of infection and the inter-host exchange of small molecules. In all cases, the levels of fate-influencing molecules (Als or arbitrium) are connected to the population context. However, it is not well understood which strategy would be optimal given variation in environmental conditions.

A longstanding rationale for the adaptive benefit of lysogeny is that integration by phage into their bacterial hosts enables viruses to persist even when fluctuations drive bacterial densities to low levels. Using a nonlinear model of population dynamics between phage and bacteria, Stewart and Levin [23] found that temperate phage densities were higher than that of virulent phage densities following transient periods of poor conditions for bacterial survival. In the context of a widespread lytic infection, as the susceptible population reduces, the benefit of lysis diminishes. Thus, in the early stages of infection, higher virulence may be selected and as bacterial populations decrease, more temperate phage are successful [5], allowing the bacterial population to recover. A putative inverse relation between susceptible population density and the benefit of lysogenic pathways has been suggested using evolutionary models [24] as well as in conceptual models of stochastic growth [3, 18]. By extension, if an infecting phage could estimate bacterial cell densities or the fraction of infected cells, then there is a potential adaptive benefit for the evolution of a decision switch that could modulate the rate of initiating lysis or lysogeny.

Here, we focus on phage of the SPbeta group and the arbitrium system to explore the potential adaptive benefits of arbitrium-dependent decision switches. In doing so, we utilize a control-theoretic framework embedded as part of a nonlinear population model to identify optimal responses of phage to variation in arbitrium concentration. We select a time-horizon to maximize near-term fitness, as measured in terms of the increase in infected cells – which correspond to epidemiological birth states for phage. This allows us to explore the link between strategies and fitness given variation in near-term selection horizons. As we show, switching between lysis to lysogeny with increasing arbitrium represents a control-theoretic optimum to maximize fitness in the near-term. In the Discussion we consider ways to extend the formalism to evaluate long-term adaptation and speculate on how the evolution of communication-based systems can help shape long-term phage strategies.

## 2 Methods

### 2.1 Population dynamics model of virus-bacteria-arbitrium system

We consider the population dynamics of temperate phage in a resource-explicit model that includes explicit representation of infections including cells that are either susceptible (*S*), exposed (*E*), lytic-fated infected (*I*), or lysogens (*L*), as well as virus particles (*V*), see Fig. 1. The exposed cells have been infected but denote the situation in which the virus has not yet committed to either the lytic pathway (turning it into an lytic-fated infected, *I*, cell) or the lysogenic pathway (turning it into a lysogen, *L*). The lysis-lysogeny decisions are guided by the concentration of a small quorum sensing peptide *A* (i.e., arbitrium) [11]. For the most general case, we assume arbitrium molecules are produced from lytic-fated infected cells and lysogens at some (possibly different) generation rate. Here, we denote the probability of lysogeny as a response function of *A, i*.*e*., *P* (*A*), consequently, a single exposed cell *E* has a probability 1 − *P* (*A*) entering the *I* state. We represent this model in terms of systems of nonlinear ordinary differential equations (ODEs), which can be written as follows

**Figure 1:**
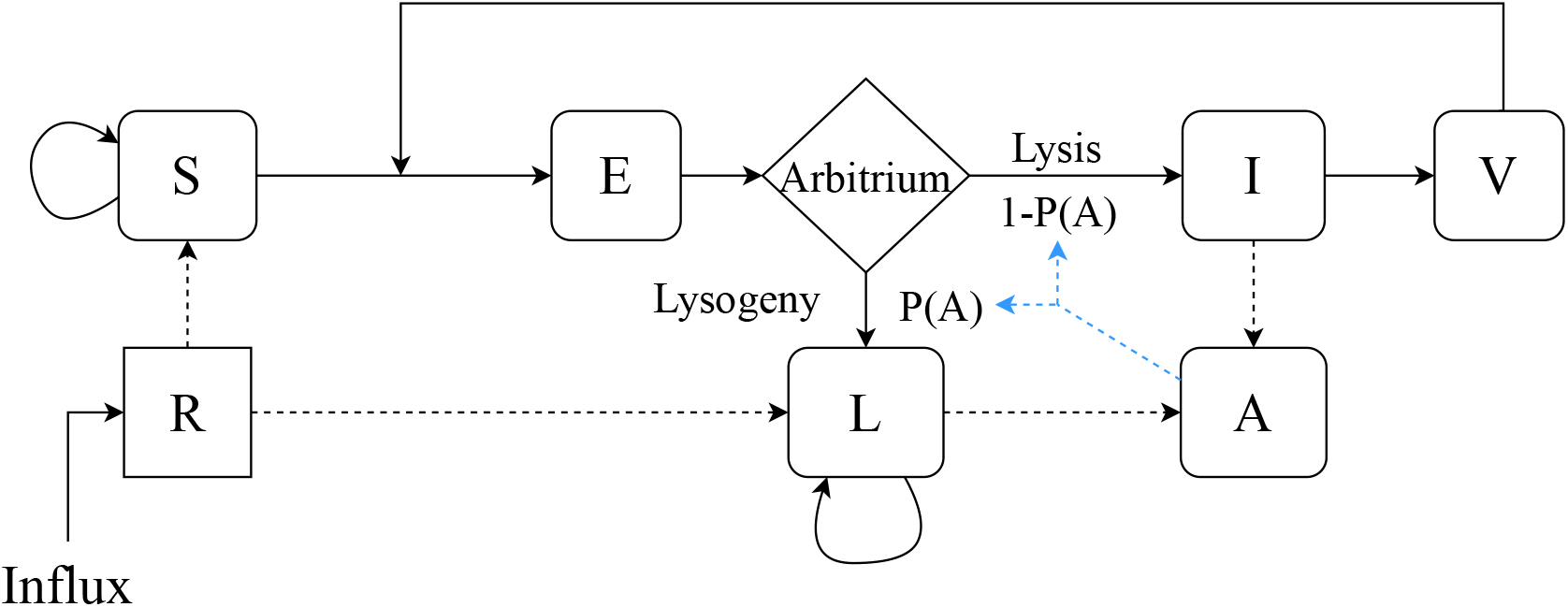
Schematic of the model. We consider a batch culture of susceptible bacteria, *S*, infected by free temperate phage, *V*, given limited resources, *R*. Early in the infection, exposed bacteria, *E*, sense the concentration of arbitrium quorum sensing molecule, *A*, from the environment and a decision between lysis or lysogeny is made based on the optimal probability function *P* (*A*). Lytic infections, *I*, result in the destruction of the infected bacteria and the release of a burst of new phage to the environment. Lysogenic infections result in the internalization of the phage in the bacterial chromosome, and the resulting lysogen, *L*, continues its replication cycle. Both infections pathways produce and release arbitrium molecules into the environment.

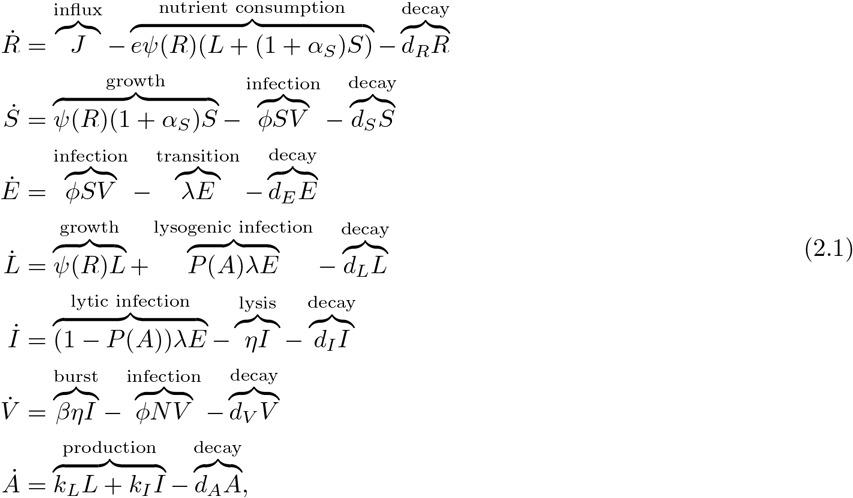

where *S, E, L, I, V* and *A* denote the densities of susceptible cells, exposed infected cells, lysogens, lytic-fated infected cells, virus particles and arbitrium molecules respectively. *N* = *S* + *E* + *L* + *I* is the cell density of all bacterial cells. Only viruses interacting with a susceptible cell lead to a new infection. Given the resource explicit dynamics, *ψ*(*R*) = *µ*_*max*_(*R/*(*R* + *R*_*in*_)) is the Monod equation, where *µ*_*max*_ and *R*_*in*_ are the maximal cellular growth rate and the half-saturation constant, *J* and *d*_*R*_ are the influx and decay rates of resources, *e* is the conversion efficiency and *α*_*S*_ is the selection coefficient that measures the relative difference in the reproductive output between lysogens and susceptible cells. Parameter *ϕ* is the adsorption rate, *d*_*S*_, *d*_*E*_, *d*_*L*_ and *d*_*I*_ are the cellular death rates of susceptible cells, exposed infected cells, lysogens and lytic-fated infected cells respectively. In addition, *d*_*V*_ and *d*_*A*_ are the decay rate of viruses and molecules. *λ* is the transition rate from exposed cells to the fate determined cells, *η* is the lysis rate, *β* is the burst size. The arbitrium molecules are produced by *L* and *I* at a production rate of *k*_*L*_ and *k*_*I*_ respectively. According to [11], the binding of the arbitrium molecule and the *AimR* receptor of phage phi3T saturates around 500 nM (3 × 10^14^ molecules*/*ml), which we set as the target accumulation level of arbitrium after 24 hours. These target accumulation levels are reached in our simulations when we set *k*_*L,I*_ = 5 × 10^7^ molecules*/*cell h^−1^ (∼ × 2 10^6^ molecules produced per bacterium), which fall within physiological levels of bacterial protein production [19]. For simplicity, we assume that lysogens and lytic-fated infected cells have identical arbitrium production rates. The model does not account for cellular active transport mechanisms, which could imply that lower external concentrations of arbitrium (and lower production rates) could nonetheless generate physiologically relevant internal concentrations of arbitrium. As we are focusing on near-term fitness, induction is ignored in the model (see Discussion for potential ways to extend the current model to include induction process when evaluating long-term fitness). All parameter values used are listed in Table 1. For tractability, we assume the probability function *P*_*θ*_: ℝ^+^ → [0, 1] lies in the class of sigmoid functions parameterized by *θ*, where *θ* = [*k, A*_0_]^*T*^; and contains shaping parameter, *k* ∈ ℝ, and the switching point, *A*_0_ ∈ ℝ^+^. In doing so, the probability of lysogeny has the form

**Table 1:** Parameter choices from previous works [26]. †, ‡:*k*_*L,I*_ estimation from data in [11] (see 2).

| Parameter | Value | Unit |
| --- | --- | --- |
| $e$, conversion efficiency | $5 \times 10^{-7}$ | $\mu g$ |
| $\mu_{max}$, maximum growth rate of cells | 1.2 | $h^{-1}$ |
| $R_{in}$, half saturation constant | 4 | $\mu g/ml$ |
| $\lambda$, transition rate from exposed cells | 2 | $h^{-1}$ |
| $\eta$, lysis rate | 1 | $h^{-1}$ |
| $J$, influx rate of resources | 0 | $\mu g/ml h^{-1}$ |
| $d_R$, decay rate of resources | 0 | $h^{-1}$ |
| $d_S$, cellular death rate of susceptible cells | 0.075 | $h^{-1}$ |
| $d_E$, cellular death rate of exposed cells | 0.075 | $h^{-1}$ |
| $d_L$, cellular death rate of lysogens | 0.075 | $h^{-1}$ |
| $d_I$, cellular death rate of lytic cells | 0.075 | $h^{-1}$ |
| $\beta$, burst size | 50 | |
| $\phi$, adsorption rate | $3.4 \times 10^{-10}$ | $ml/h$ |
| $d_V$, virion decay rate | 0.075 | $h^{-1}$ |
| $\alpha_S$, selection coefficient | 0.1 | |
| $k_L$, generation rate of arbitrium molecules from lysogens | $5 \times 10^{7\dagger}$ | molecules/cell $h^{-1}$ |
| $k_I$, generation rate of arbitrium molecules from lytic cells | $5 \times 10^{7\dagger}$ | molecules/cell $h^{-1}$ |
| $d_A$, decay rate of arbitrium molecules | 0.1 | $h^{-1}$ |

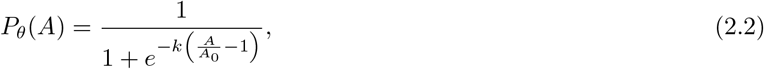

which is a function of the arbitrium molecule concentration. Note that the Model 2.1 uses a nonlinear population modeling approach (similar to Doekes et al. [10]), but differs from [10] in key ways: (i) includes greater specificity of the intracellular decision process; (ii) includes the explicit potential for lysogens to produce arbitrium.

### 2.2 Near term fitness and growth rate

We quantify viral fitness in terms of the reproduction of infected cells in order to compare pure lytic and context-dependent lysis-lysogenic strategies. Malthusian fitness is expressed as the growth rate of phage-infected cells, from either the lytic or lysogenic pathway. Newly infected cells can only be produced by the epidemiological birth states, i.e. the lysogens and exposed cells [25, 16]. The exponential growth rate *ρ*, denotes the effective reproductive rate of these infected cells and is given as:

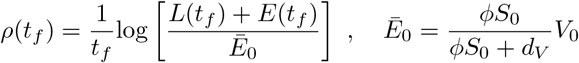

where *Ē*_0_ is the average number of cells infected in the first generation, the first factor of *Ē*_0_ denotes the probability that a free virus particle is adsorbed into a susceptible cell before it decays. For a given time horizon, the fitness monotonically depends on the sum of the lysogens and exposed cells produced at the final time relative to the initial density of exposed cells.

### 2.3 Optimization framework for fitness maximization through probability function selection

We formulate an optimization problem over different probability functions with the aim of achieving a maxima of the near term fitness, as defined in the previous section. By controlling the lysis-lysogeny decisions *p*_*θ*_(*A*) (more precisely, the variable *θ*), the objective is to maximize the the epidemiological birth states at the final time. In order to do so, we define a payoff as a function of *θ, J*(*θ*) = *L*(*t*_*f*_) + *E*(*t*_*f*_), such that the optimization problem can be written as

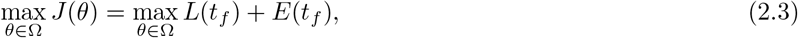

where Ω = ℝ × ℝ^+^.

We implement a gradient descent algorithm to numerically compute the optimal *θ* for optimization problem (2.3). In order to do so, we need to evaluate the gradient of *J* w.r.t *θ*. Let the state vector of the system be represented by *X* ∈ ℝ^*n*^, (here *n* = 7) and let the dependence of the payoff function on the final state be given by Ψ, *i*.*e*., *J* = Ψ(*X*(*t*_*f*_)) = (*e*_3_ + *e*_4_)^*T*^ *X*(*t*_*f*_) as J is linear. Here *e*_*i*_ represents the *i*^th^ unit vector. We represent the nonlinear system (2.1) as

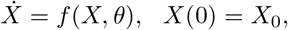

where *f* (*X, θ*) ∈ ℝ^*n*^, is the right-hand-side (RHS) of system (2.1).

Note that *J* is a function of the final state, *X*(*t*_*f*_), where *X*(*t*_*f*_) is a function of *θ*, thus we apply chain rule for finding the required derivative. In doing so, the derivative of *J* with respect to *θ* takes the form of

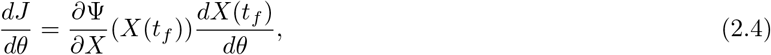

where 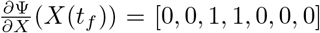 is a row vector. There is no closed-form algebraic expression for the terminal state derivative, *i*.*e*.,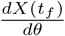. To evaluate it, we let 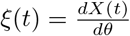, and using 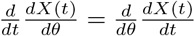, we have

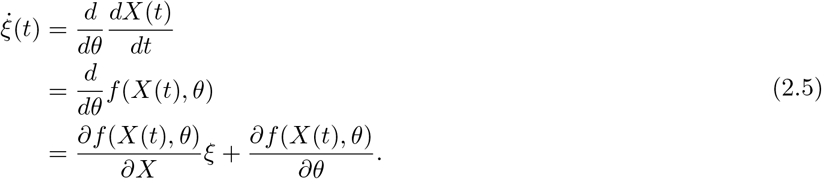

Note that system (2.5) is a linear system. Let the state transition matrix for (2.5) be Φ. Then, we obtain

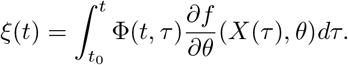

Substituting the terminal state derivative *ξ*(*t*_*f*_) in to Eq. (2.4), the derivative of *J* can be written as

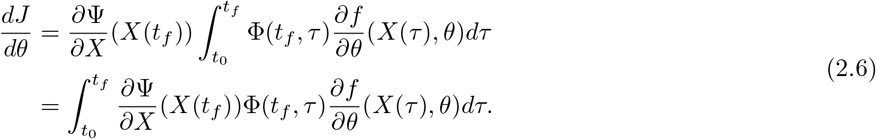

Here, we define the *costate* as 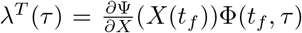, with 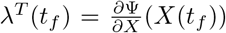. Differentiating the costate, we have

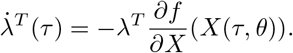

The costate trajectory *λ*(*t*) can be solved numerically in backward direction using the terminal boundary condition. Then, we substitute the costate in Eq. (2.6) and obtain

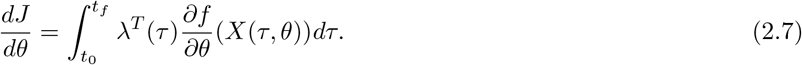

Since the costate trajectory can be computed by numerical integration of the costate differential equation, the derivative of *J* w.r.t. *θ* can be computed.

The gradient descent algorithm does not guarantee the convergence to the global maxima. To circumvent the issue of convergence to a local maxima, we perform ‘naive sampling’ before the gradient descent search. First, we divide the search space into grids and find the payoff function’s value at the center of each grid. Then, we select the center-point which gives the largest value for *J* and choose that as the starting point of the gradient descent algorithm. If the grids are small enough, this naive sampling approach will be equivalent to a ‘brute-force search’ of the complete parameter space. Here, we employ Armijo step-size in gradient descent [2]. This process is iterated until the increase in cost function between iterations is lower than the preset termination value. Once the optimal parameters are reached, they can be used to obtain the optimal probability function.

### 2.4 Arbitrium sensing mechanism and expected lysogeny probability

The inclusion of imperfect sensing is modeled via a sigmoidal response function that is dependent on the estimated concentration instead of the true concentration. Let the concentration of arbitrium molecule in the medium be *A* molecules/ml. We assume a typical bacterial size of ~1 *µm*^3^. As such, the number of molecules per cell volume would be on the order of *A/*10^12^ (assuming nearly uniform concentration in the medium), and the estimated value will be denoted as *Â*.

Assume there are *E* exposed cells, numbered as 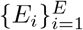 medium be *A* and let the sensed concentration be 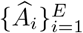. Let the true concentration of arbitrium molecule in the for each phage in an exposed cell. For convenience, we define 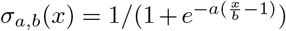 as a standard sigmoid function parameterized by *a* and *b*. In doing so, we can write the probability of lysogeny 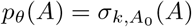. The probability of lysogeny for exposed cell *E*_*i*_ can be written as

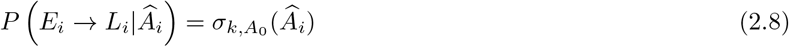

as it only depends on the sensed concentration. Note that *A*_0_ is the switching point for the sigmoid and *k* is a shaping parameter, which are the two parameters to be optimized.

For each individual cell with a sensed concentration of *Â*_*i*_, we denote its initiation of lysogeny with an indicator function, 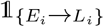, a binary random variable. Then, the total number of exposed cells that undergo lysogeny, denoted by *L*_new_, can be written as

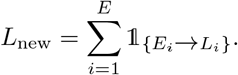

The conditional expectation of *L*_new_ given the estimated concentrations 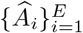 is

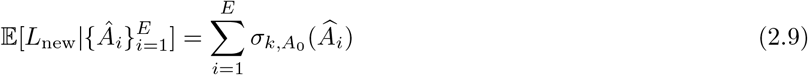

Assuming *Â*_1_, *Â*_2_, …, *Â*_*E*_ are *i*.*i*.*d* Poisson random variables, using the iterated expectation law on (2.9), we have

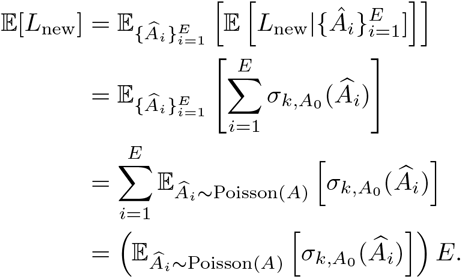

We approximate the first moment of the lysis-lysogeny decision function of the estimated concentration, *i*.*e*., 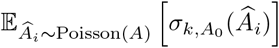, using Taylor expansions provided that 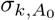 is sufficiently smooth (twice differentiable at least) and the moments of *Â*_*i*_ are finite. Taking the local expansion of 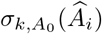 around *A* yields

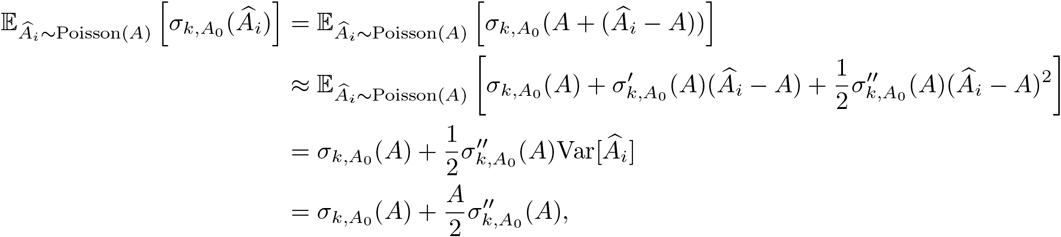

where the first term corresponds to the mean lysis-lysogeny response of estimated concentration and the second term corresponds to the *propagation of uncertainty*. Hence, we approximate the expectation by the first three terms from the Taylor series. Then, we can compute the expected number of exposed cells that undergo lysogeny as

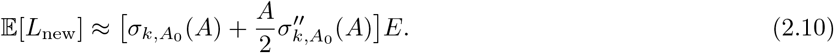

The subsequent results in all sections account for sensing noise and use (2.10) for estimating the lysogeny probability.

### 2.5 Simulation Details

The parameter values used for these results are given in Table 1. The initial concentrations of susceptible and viral particles are 10^7^ cells/ml and 10^5^ phage/ml respectively (MOI=0.01) and the rest are set to 0. The initial resource concentration is 40 *µg/ml* with no influx, except when the effect of resource level on switching point is studied (where the values used are explicitly stated in text). Simulation code is written in MATLAB R2020a and available for download at https://github.com/WeitzGroup/LysisLysogeny.

## 3 Results

### 3.1 Analysis of fixed probability decision strategies

We begin by evaluating the population dynamics of host-phage systems in which phage have a fixed probability to initiate lysogeny, *P*. We first consider three cases - that of purely lytic (*P* = 0), mixed (*P* = 0.5), and purely lysogenic (*P* = 1) phage (first, second and third panels of Figure 2a). The system is initialized with a low resource density, an MOI of 0.01, where MOI is the initial virus-host ratio, and then simulated for 48 hours.

**Figure 2:**
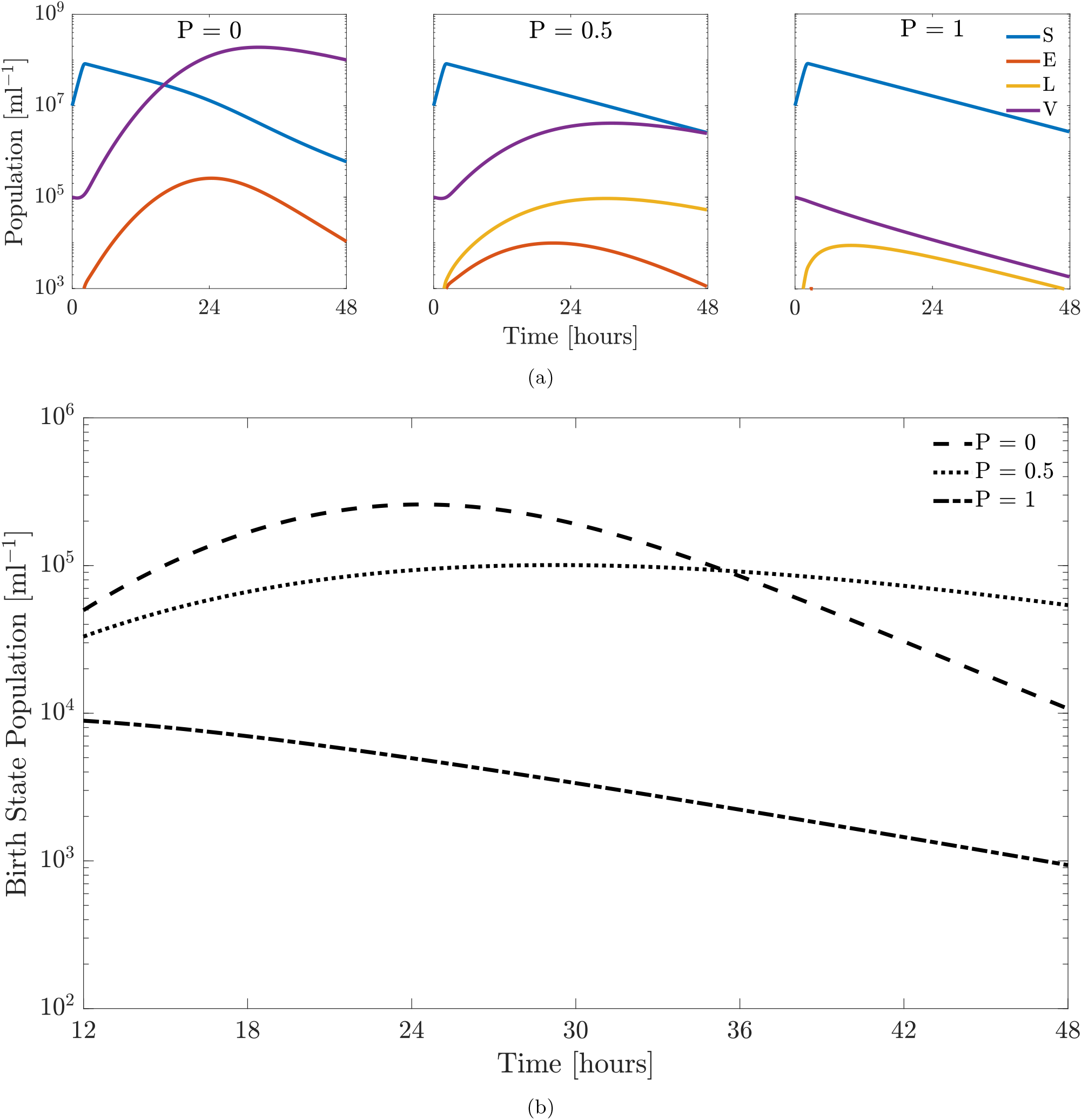
Temperate phage-bacteria infection dynamics for different fixed probabilities of lysogeny (*P* = 0, *P* = 0.5 or *P* = 1 where *P* is the probability of lysogeny) for 48 hours with an MOI of 0.01. The model parameters are given in Table 1. **(a)** Population dynamics for the system when a fixed strategy is employed over a period of 48 hours. The three panels correspond to an obligately lytic (*P* = 0, left panel), stochastic (*P* = 0.5, middle panel) and purely lysogenic (*P* = 1, right panel) strategy, respectively. **(b)** Comparison of total birth states, i.e. total population of lysogens and exposed cells, for the three fixed strategies. The optimal fixed strategies with highest birth states population vary with time. Specifically, obligately lytic strategy is favored for relative short time scales (from 12 hours to 36 hours). Beyond this, the mixed strategy (*P* = 0.5) is favored.

The first, second and third panels of Figure 2a show the dynamics of the system if the decision switch is obligately lytic (*P* = 0), stochastic with an equal probability of lysis or lysogeny (*P* = 0.5) or in which the phage always initiate lysogeny (*P* = 1). In each case, the susceptible cells show a transient increase due to the initial resource concentration, followed by a decline as a consequence of resource scarcity and cellular decay. Lysis also contributes to the decline for the cases of *P* = 0 and *P* = 0.5. In the first case of obligate lysis, the viral population quickly increases and a large number of exposed cells are generated, without the formation of lysogens. In the second case, the viral population does not increase as quickly as in the case of a purely lytic strategy due to the fact that half of the infections generate lysogens (which do not produce more virus particles). The mixed strategy also leads to lysogens (in addition to exposed cells), both of which are considered epidemiological birth states. For the last case of pure lysogeny, there is no increase in the virus particles density given the absence of lysis in the initial decision switch (as noted, the model does not include an induction process). This means the virion density rapidly decreases due to decay and infection of host cells. A purely lysogenic strategy does generate both types of birth states, i.e. exposed and lysogens, however the low MOI results in very few bacterial cells getting infected. Consequently, the total aggregate density of exposed cells and lysogens is significantly less than that of either the purely lytic or stochastic strategy.

We compare the relative fitness of the purely lytic (*P* = 0), mixed (*P* = 0.5) and purely lysogenic (*P* = 1) cases over a 48 hr time horizon. As stated previously, the cell-centric metric of fitness depends on the population of lysogens and exposed cells. Fig. 2b compares the birth state population for all three strategies over the time horizon of 12 to 48 hours. Initially, from 12 to 36 hrs, obligate lysis appears to be the dominant strategy. The low MOI, coupled with the short time horizon prevents a host population collapse and a large number of exposed cells are generated through lysis. However, for long time scales, i.e. beyond 36 hours, a pure lytic strategy depletes the bacterial population such that the total number of exposed cells begins to decline and the mixed strategy (*P* = 0.5) becomes more favorable.

### 3.2 Effect of time horizon on optimal strategies for fitness maximization

We evaluate the near-term fitness of temperate phage using a communication molecule for lysis-lysogeny decisions and show how time horizons shape the responsiveness to the signaling molecule. Hence, rather than using a fixed value of *P*, the infected cells are lysogenized as a function *P* (*A*) where *A* is the concentration of arbitrium in the system. We initialized the system with 10^7^ bacteria and a MOI of 0.01. For each time horizon we use the optimization framework described in 2.3 to calculate the optimal response function to arbitrium molecule – optimal probability function *P*_*opt*_(*A*) – that maximizes near-term fitness of the phage population.

Figs. 3a and 3b show the optimal probability functions for all time horizons between 12 to 48 hrs, *P*_*opt*_(*A* |*T*_*f*_), with arbitrium production rates of 5 × 10^7^ molecules*/*cell h^−1^ and 10^7^ molecules*/*cell h^−1^ respectively. For the different generation rates, the switching strategy is robust and a corresponding shift in the switching point (from lysis to lysogeny) can be observed.The switching point scales linearly with the production rate of arbitrium (see Fig. S1), maintaining a pattern of an initial lytic period followed by a lysogenic period in the phage dynamics. Therefore, we set the arbitrium production rate to be 5 × 10^7^ molecules*/*cell h^−1^ for subsequent analyses, recognizing that changes in production rate will also shift the expected switching point as a function of *A*.

**Figure 3:**
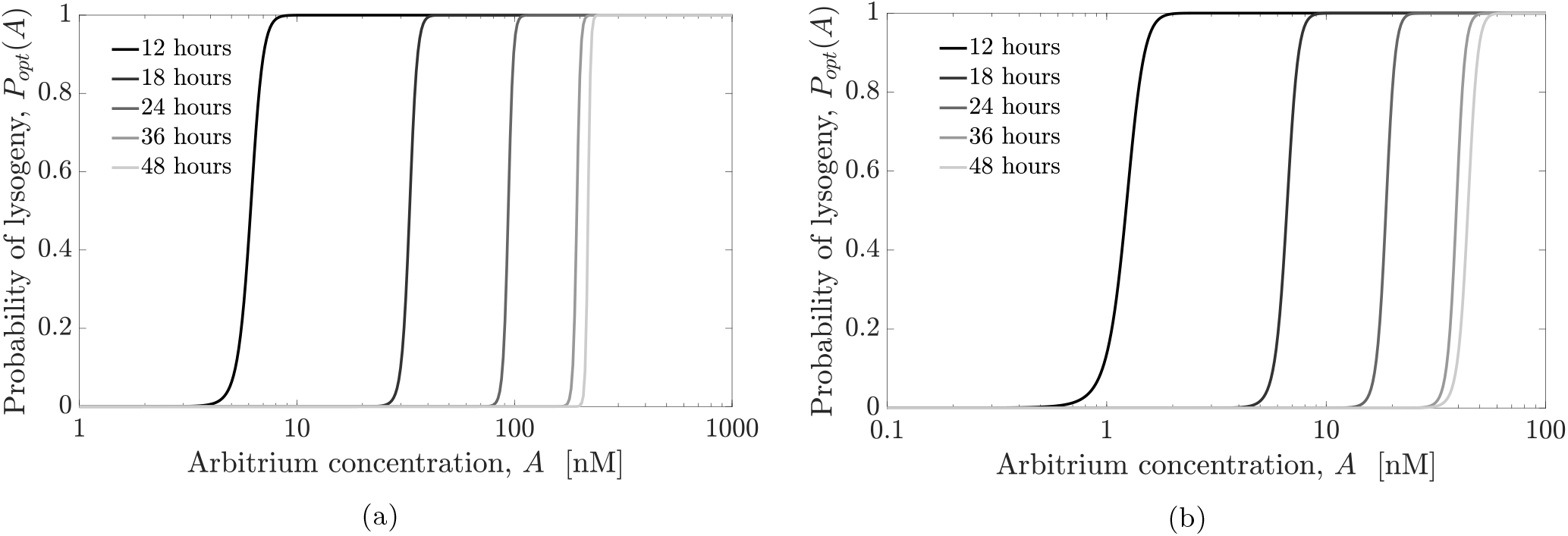
Comparison of the optimal lysis-lysogeny decision response functions (function of arbitrium molecule concentration) given variations in time horizon (from 12 hours to 48 hours). The switching point (from lysis to lysogeny, i.e., from *P* = 0 to *P* = 1) in the sigmoidal lysis-lysogeny response function shifts to the right as the horizon increases. **(a)** Production rate of arbitrium molecules used is 5 × 10^7^ molecules*/*cell h^−1^ as given in Table 1. **(b)** A lower production rate of 10^7^ molecules*/*cell h^−1^ is used in this case, which results in a corresponding shift of optimal strategies to lower concentrations.

For each function, the probability of lysogeny is low when the concentration of arbitrium molecule is small and the probability of lysogeny is high when the concentration of arbitrium is high. The switch from lysis to lysogeny is always sharp for every optimal probability function, which indicates that sensitivity to arbitrium is high upon the switch point. The switching points of the optimal probability functions increase with longer time-horizons, which indicate phage might decrease their responsiveness to arbitrium for longer time-horizons. We note that for time horizons from 12 to 24 hours, the probability of lysogeny was lower than 0.5 for approximately 80% of the total period, while for 48 hours, it was close to 50% of the period. The two observations together mean that for longer time horizons, a larger initial period of lysis followed by a switch to lysogeny is optimal. However, the fraction of time for which lysis occurs decreases with increasing time horizon. Another notable trend is that as the time horizon increases uniformly, the gap between the switching points decreases which suggests that the switching points would not increase indefinitely. The observations of the fractional period of lysogeny increasing with the time horizon and reduction in the gaps between the optimal probability functions show that with increases in the time horizon, the optimal strategy places more weight towards generating lysogens, which are longer lived than exposed cells.

### 3.3 Population Dynamics for Optimal Switching Strategies

We now examine the population dynamics of the full system given optimal switching strategies. Fig 4 shows the population dynamics for the susceptible and exposed cells, lysogens and virus particles for time horizons of 12, 18, 24, 36 and 48 hours. From the dynamics for all time horizons, we can observe common trends. The susceptible population increases due to the initial resource concentration followed by a decline due to resource depletion, cell decay and lysis. Next, the concentration of arbitrium molecule takes time to build up in the system, thus the phage are predominantly lytic at the start. Further, the MOI is low, thus there is an abundance of susceptible cells in the medium. These factors explain the rapid growth of viral particles at the start of the interval and consequently, a large population of exposed cells is generated. The end of the initial period of lysis can be observed from the point where the population of lysogens starts increasing. This signals a switch from a predominantly lytic strategy to a lysogenic strategy. In this phase, the viral particle population declines (due to decay and integration with host cells) and the lysogens increase. As noted previously, an increase in the time horizon delays the switch to lysogeny. However, the fraction of time with a predominantly lysogenic strategy increases with the time horizon.

**Figure 4:**
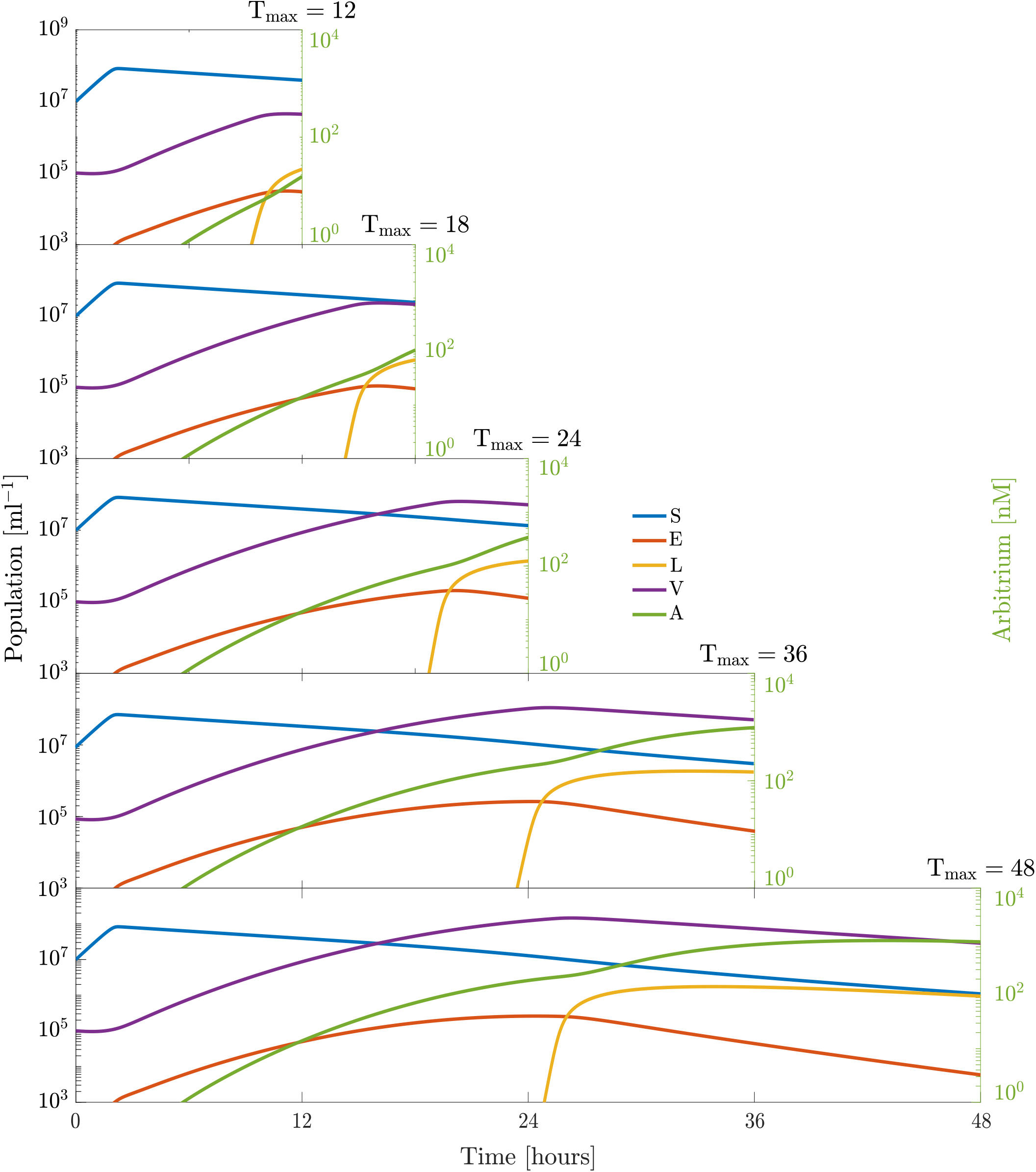
Temperate phage-bacterial infection dynamics for an optimal switching strategy in the lysis-lysogeny decision as a function of the final time horizon. Consecutive panels show an optimal strategy dynamics resulting from the optimal response functions to arbitrium quorum sensing concentration represented in Fig. 3a given a time horizon (12, 18, 24, 36 and 48 hours, respectively) We see a common strategy of pure lysis followed by a stochastic strategy in all cases. The time of the switch from obligate lysis to a stochastic strategy can be deduced by the time at which the production of lysogens start. The switch occurs based on the length of the time horizon, with a later switch for longer horizons. The initial MOI is set as 0.01, the additional model parameters are given in Table 1.

### 3.4 Near-term fitness benefits of optimal switching strategies

In order to evaluate the fitness benefits of lysis-lysogeny switching based on responses to communication signals, we compare the fitness of phage populations using an optimal strategy with that resulting from fixed strategies. Fig. 5a shows the fitness values for the optimal probability strategy for 48 hours and different fixed probability strategies (*P* = 0, *P* = 0.1, …, *P* = 0.9, and *P* = 1). The strategy based on communication performs better than any fixed probability strategy in terms of the sum of the lysogens and exposed cells at the final time. Optimal switching is more than 4 times better than the highest value achieved for a fixed strategy (for a fixed probability of 0.2), which demonstrates that a phage following such a communication strategy would have a higher near-term fitness compared to a phage following any other fixed probability strategy.

**Figure 5:**
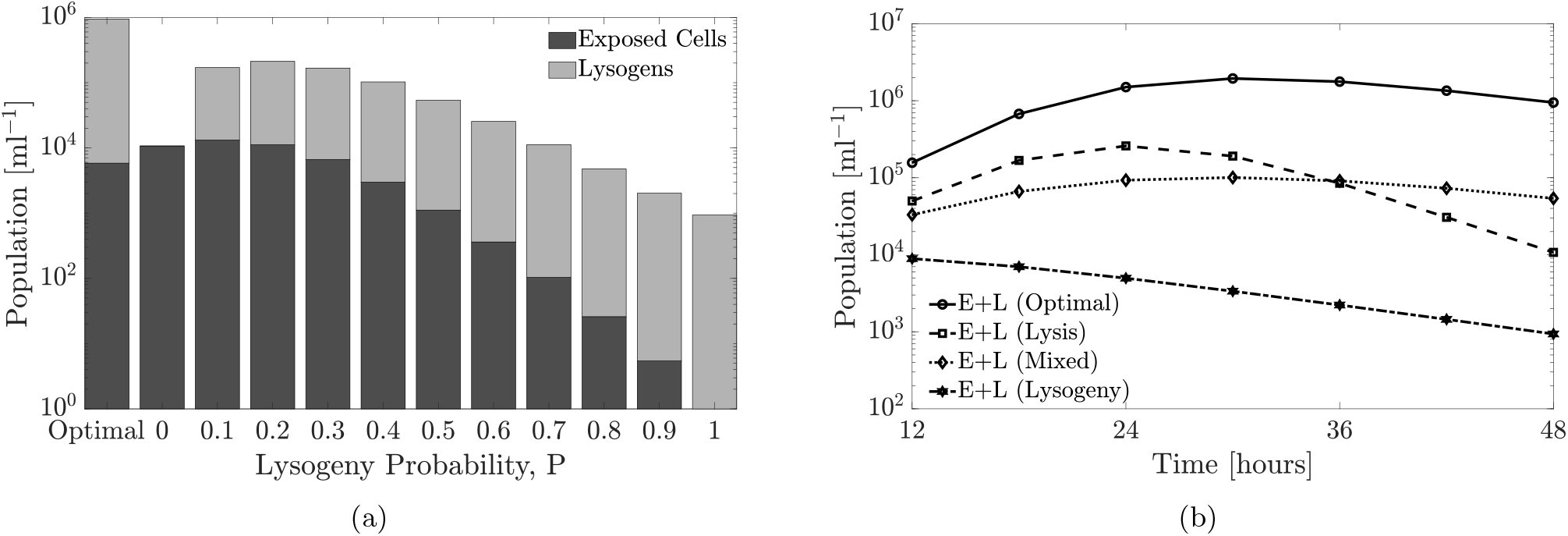
Phage fitness comparison for optimal and fixed probabilities of lysogeny. The relevant parameters are present in Table 1. Note that the optimal strategy is a response function of arbitrium molecule concentration. The initial MOI is set as 0.01, the additional model parameters are given in Table 1. **(a)** Comparison between birth states produced from multiple fixed and optimal probability strategies for a time horizon of 48 hours with an MOI of 0.01. Here, we consider ten alternative fixed probability strategies for lysis-lysogeny i.e. not reliant on arbitrium molecule concentration. We simulate with probabilities from 0 to 1 with increments of 0.1 and compare the performance of all these strategies to the optimal strategy which utilizes the arbitrium system. The comparison shows the total birth states produced by the optimal strategy is higher than all fixed strategies. **(b)** The final time birth states population vary with time horizon (from 12 hours to 48 hours) and strategies (optimal, pure lysis, pure lysogeny and stochastic). The comparison is made for all time horizons from 12 to 48 hours, but only three fixed strategies are considered for clarity. The birth states formed by following the optimal strategy are greater than these fixed strategies for all time horizons.

Next, we perform the same analysis for all optimal strategies for the time horizons of 12 to 48 hours and compare them to three fixed strategies (*P* = 0, *P* = 0.5, and *P* = 1). Fig. 5b presents the total birth states for all time horizons (12, 18, 24, 30, 36, 42 and 48 hours) and compares them to the birth states produced by the fixed strategies. The corresponding optimal strategy produces more birth states compared to any fixed strategy for all time horizons, providing evidence that communication-based switches from lysis to lysogeny may be of adaptive benefit to temperate phage.

We recognize that measuring total birth states depends on the sum of lysogens and exposed cells, irrespective of composition. To explore shifts in the contribution to fitness, we examined the fraction of lysogens and exposed cells at the final time. As the time horizon increases, the fraction of lysogens monotonically increases from 0.82 at 12 hours and approaches 1 at 48 hours, with the corresponding reduction in the fraction of exposed cells. This implies that given sufficiently long interaction times, optimal switching strategies predominantly rely on lysogens as the basis for increasing near-term fitness. On the cellular level, as stated above, this occurs due to a build-up of arbitrium molecule in the medium over time, leading to inhibition of lysis and switching to a strategy dominated by lysogeny. However, in order to increase lysogens at the final time, the strategy initially tends toward lytic outcomes. Lysis depletes susceptible cells, which reduces niche competition between them and increases the potential benefits of vertical transmission.

### 3.5 Effect of resource level on optimal strategy

Thus far we have evaluated changes in adaptive switching strategies on near-term fitness given an environment without influx of resources for cellular growth, analogous to batch culture conditions. Next, we evaluate the impact on different resource conditions, including (i) increased initial concentration of resources and (ii) resource influx which maintains higher levels of resources in the environment throughout the focal period similar to chemostat growth conditions. As before, we use the optimization approach described in the 2.3 to identify an optimal strategy, including an optimal switching point, *A*_*c*_ where the system tends to switch from lysis to lysogeny.

Fig. 6a shows three heatmaps plots of the optimal strategy for different time horizons and initial resource concentrations. In all three panels the time horizon varies from 12 to 48 hours, and the initial resource concentration starts from a base value of 40 *µg/ml* up to 100 *µg/ml*. The left panel shows the optimal switching point for a given time horizon and initial resource concentration. For any particular time horizon, as the initial resource level increases, the switching point also increases implying a longer period of a lytic-like strategy. This delay in switching occurs due to the susceptible population rising to a higher value (due to the greater initial resource concentration). However, we also note that the the switching point need not inevitably increase for longer time horizons, *T*_*f*_ (see cases of initial resource concentration of 80 and 100 *µg/ml* in Fig. 6a). This discrepancy can be explained as follows. For these two initial concentrations, the susceptible population rapidly increases due to the abundance of resources. Given sufficiently short time horizons, phage can maximize fitness by preferentially producing more lytically-infected cells, which imply a higher switching point. This strategy of investing in lytically-infected cells works in the limit of higher resource concentration, which prevents a population collapse for shorter periods. For longer periods, we again see a return to a lower switching point, which would translate to investing in the production of lysogens. The fraction of lysogens (middle panel) increases with an increase in time horizon, as noted in the previous section. This trend is true for all initial resource concentration values. We observe a monotonically increase of the growth rate of phage with the initial resource level (right panel Fig. 6a). However, as time horizon increases, the growth rate is observed to decrease, i.e. the short-term fitness decreases with an increasing time horizon. The reduction in fitness as a result of an increase in time horizon, further suggests that environmental factors that alter host populations, such as nutrient depletion may drive selection of communicating phage to become more sensitive to arbitrium concentration when conditions are less favorable for the host. The initiation of lysogeny when hosts are scarce increases the chances of increasing infected cells, albeit in an integrated form.

**Figure 6:**
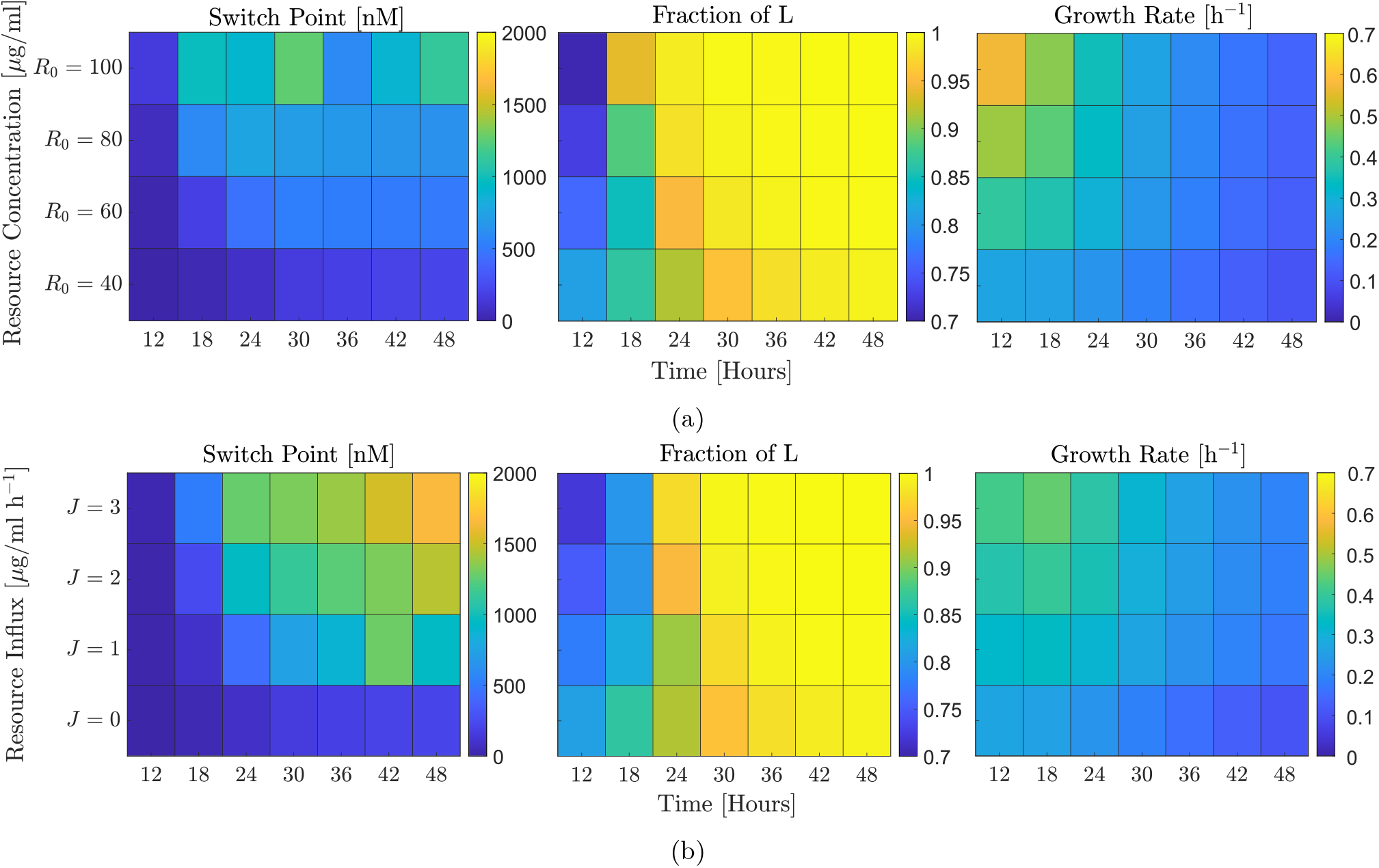
Effect of resource level (change in initial concentration and influx) on optimal lysis-lysogeny switching point. The simulation parameters are present in Table 1. **(a)** Heatmaps for time horizons from 12 hours to 48 hours and variation in initial resource concentration, from 40 *µg/ml* (which is the base case concentration used in all previous plots) to 100 *µg/ml*. The three panels correspond to the optimal switching point (left panel), fraction of lysogens at the final time (middle panel) and growth rate (right panel). **(b)** Heatmaps for time horizons from 12 hours to 48 hours and variation in resource influx, from 0 *µg/ml h*^−1^ (which is the base case concentration used in all previous plots) to 3 *µg/ml h*^−1^. The three panels correspond to the optimal switching point (left panel), fraction of lysogens at the final time (middle panel) and growth rate (right panel).

Fig. 6b shows three heatmap plots of the optimal strategy for different time horizons and resource influx. In all three panels the time horizon varies from 12 to 48 hours, and the resource influx varies from 0 (no influx) to 3 *µg/ml h*^−1^. The left panel shows the optimal switching point for a given time horizon and resource influx. For any particular time horizon, increasing the resource influx also increases the switching point, as before. Further, the switching point concentration of arbitrium molecule strictly increases with increasing time horizon for the considered values of resource influx. The susceptible population does not increase as rapidly, which we interpret to mean that that the optimal strategy is for phage to invest in lysogens. The fraction of lysogens (middle panel) also follows a similar increasing trend with an increase in time horizon, and is true for all resource influx values. Finally, the right panel shows the growth rate. Again, a higher influx translates to a higher growth rate. Similar to the trend observed before, the growth rate decreases with an increase in time horizon, i.e. the short-term fitness decreases with an increasing time horizon.

## 4 Discussion

We have developed an optimal control framework based on the SPbeta phage arbitrium system to study the near-term adaptive benefits of responsive switching between lysis and lysogeny. We used a cell-centric metric to compare temperate phage fitness in the near-term. Optimal strategies computed through this technique all share the common feature of a high lysis probability when the concentration of arbitrium molecules are low and a high lysogenic probability when the concentration of arbitrium molecules are high. Short horizons favor a primarily lysogenic strategy because the increase in viral particles through lysis does not directly lead to more infected cells at the final time. The longer the horizon, the later the switch occurs, allowing for an increase in infected cells. Thus, for temperate phage, an initial period of lysis allows sufficient growth to infect a large number of bacterial cells. After this, a switch to lysogeny ensures that the host is not depleted and the phage generated a large number of infected cells in the form of lysogens. Similarly, a higher resource concentration, either through initial concentration or influx results in comparatively longer periods of lysis followed by lysogeny. This exploits the increase in bacterial population due to higher resource concentration which can support a larger viral population without collapsing. The switching point is expected to increase in systems with more resources and/or those that can support sustained periods of host growth.

In exploring optimal adaptive strategies, we find that the time horizon plays a role in another emergent property: the relevance of lysogeny. For longer time horizons, the fraction of lysogens present at the final time increases. This occurs due to a buildup of arbitrium molecules in the medium, which increases the probability that infected cells initiate the lysogenic pathway. In this framework, we interpret the preferential use of lysis as a short term strategy for immediate increase in phage particles, and lysogeny as part of a longer time strategy to maximize the growth of the infected cell population (which we use as a metric for near-term viral fitness). Though the growth rate declines for long time horizons, lysogeny ensures that the phage can persist despite a depleted host population, which is not possible if lysis is continued indefinitely.

Even with this simplified model, we observe interesting properties which can shed light on the balance between the lysis and lysogeny pathways in a biological system. In our model, the fitness for phage that utilized a fixed strategy of lysis, lysogeny, or stochastic (but non-responsive) switching was lower than that of phage with optimally responsive switching strategies. Thus, there is a fitness advantage conferred by temperate phage that sense and ‘choose’ the optimal life cycle (lytic or lysogenic) (consistent with the findings of [10], albeit here we provide evidence in support of sharp switching, rather than assuming such switching is sharp a priori). Notably, these findings hold for resource-limited conditions, but increases in resource availability can shift the optimal strategy towards lytic outcomes (especially in the short-term).

The optimization approach used here has certain limitations. The optimal control method identifies the switching strategy with the highest near-term fitness given a fixed time horizon. However, there is no guarantee that this strategy is an evolutionary stable strategy over the long term (in related work, Doekes et al. used an eco-evolutionary modeling framework to explore the potential evolution of fixed decision strategies [10]). Likewise, due to our focus on adaptive benefits of switching to short-term fitness, we do not consider induction of lysogens in the model. The induction of prophage leading to the reinitiation of the lytic pathway would become more relevant in the long-term, particularly in stressed cells. Moreover, if the initial MOI is very low or the environment is resource-rich, then a pure lytic strategy (rather than a switching strategy) could have higher fitness for short term horizons. Finally, the cellular sensing mechanism for arbitrium molecules is a complex process which was greatly simplified by assuming the sensed concentration follows a Poisson distribution based on the mean concentration of arbitrium molecule in the entire medium - extensions should include the intracellular uptake and regulation of decisions based on internal arbitrium ‘quotas’. Nonetheless, our finding that sharp transitions are optimal in the near-term provides additional context for experimental and modeling studies of the long-term evolution of arbitrium-based switches. The integration of a changing ecology and a sensing mechanism based on cellular mechanism would improve the applicability of the model in predicting optimal strategies for phage in real environments. In doing so, it will be critical to investigate the dependence of the optimal strategy on the multiplicity of infection.

In summary, the control theoretic framework developed here demonstrates the adaptive value of a sensing mechanism for initiating lysis or lysogeny based on environmental context informed by prior infections. In doing so, our results suggest that the early lysis of cells by temperate phage provides a mechanism for rapid expansion of virus particles that can then, later, preferentially initiate lysogeny given sensing of an extracellularly released communication peptide. When viewing fitness of viruses in terms of infected cells, such changes in strategies represent a form of feedback control by which phage both modulate and reshape their ecological context. Moving forward we hope that this near-term control theoretic framework is useful in evaluating how such sensing mechanisms may evolve over longer time horizons.

## Acknowledgments

We thank Weitz Group team members for comments and feedback. This work was supported by grants from the Army Research Office (W911NF1910384) and National Institutes of Health (1R01AI46592-01).

## Appendix

**Figure S1:**
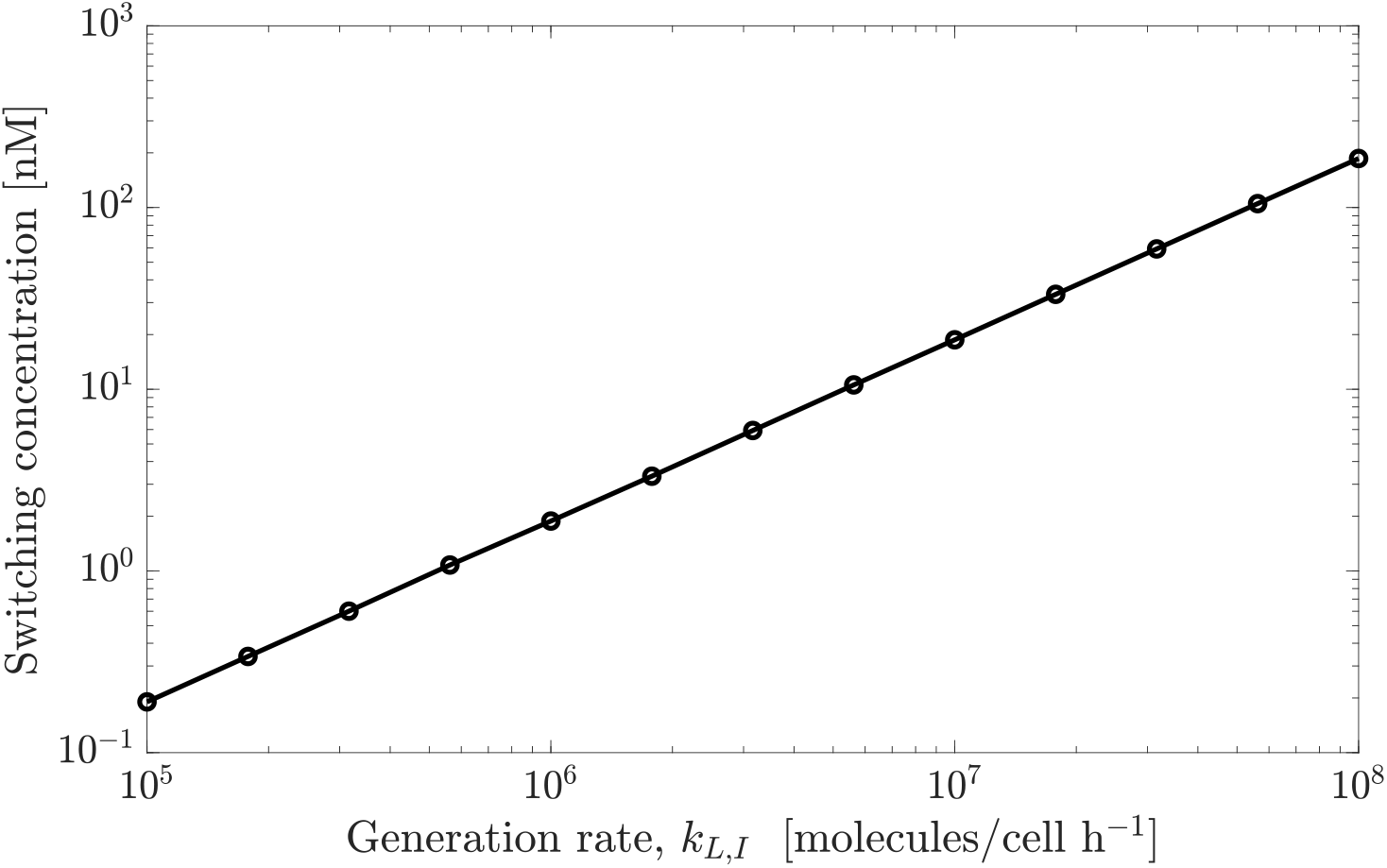
Effect of production rate of arbitrium – *k*_*L,I*_, on optimal switching concentration for a time horizon of 24 hours. The production rate is varied from 10^5^ to 10^8^ molecules*/*cell h^−1^ with exponential steps of 10^0.5^. For each arbitrium production rate, we compute the corresponding optimal switching concentration. The relationship between the optimal switching concentration and arbitrium production rate is linear (correlation coefficient = 1)

## References

[1] Stephen T. Abedon. “Bacteriophage secondary infection”. eng. In: Virologica Sinica 30.1 (Feb. 2015), pp. 3–10. issn: 1995-820X. doi: 10.1007/s12250-014-3547-2.

[2] Larry Armijo. “Minimization of functions having lipschitz continuous first partial derivatives”. In: Pacific Journal of Mathematics 16.1 (1966), pp. 1–3. issn: 00308730. doi: 10.2140/pjm.1966.16.1. url: https://www.mendeley.com/catalogue/b4a6e890-d70c-3b19-9571-5daf1a30133f/ (visited on 03/12/2021).

[3] Mikkel Avlund, Ian B Dodd, Szabolcs Semsey, Kim Sneppen, and Sandeep Krishna. “Why do phage play dice?” In: Journal of virology 83.22 (2009), pp. 11416–11420.

[4] C. Bernard, Y. Li, P. Lopez, and E. Bapteste. “Beyond arbitrium: identification of a second communication system in Bacillus phage phi3T that may regulate host defense mechanisms”. In: ISME J (Oct. 2020).

[5] Thomas W. Berngruber, Rémy Froissart, Marc Choisy, and Sylvain Gandon. “Evolution of Virulence in Emerging Epidemics”. In: PLOS Pathogens 9.3 (Mar. 2013), pp. 1–8. doi: 10.1371/journal.ppat.1003209. url: https://doi.org/10.1371/journal.ppat.1003209.

[6] Giuseppe Bertani. “Lysogeny at Mid-Twentieth Century: P1, P2, and Other Experimental Systems”. In: Journal of Bacteriology 186.3 (2004), pp. 595–600. issn: 0021-9193. doi: 10.1128/JB.186.3.595-600.2004. eprint: https://jb.asm.org/content/186/3/595.full.pdf. url: https://jb.asm.org/content/186/3/595.

[7] Joseph Bondy-Denomy, Jason Qian, Edze R. Westra, Angus Buckling, David S. Guttman, Alan R. Davidson, and Karen L. Maxwell. “Prophages mediate defense against phage infection through diverse mechanisms”. eng. In: The ISME journal 10.12 (Dec. 2016), pp. 2854–2866. issn: 1751-7370. doi: 10.1038/ismej.2016.79.

[8] A. G. Cobián Güeme, M. Youle, V. A. Cantú, B. Felts, J. Nulton, and F. Rohwer. “Viruses as Winners in the Game of Life”. In: Annual Review of Virology 3.1 (Sept. 2016), pp. 197–214.

[9] Adrienne M. S. Correa, Cristina Howard-Varona, Samantha R. Coy, Alison Buchan, Matthew B. Sullivan, and Joshua S. Weitz. “Revisiting the rules of life for viruses of microorganisms”. en. In: Nature Reviews Microbiology (Mar. 2021), pp. 1–13. issn: 1740-1534. doi: 10.1038/s41579-021-00530-x. url: https://www.nature.com/articles/s41579-021-00530-x (visited on 04/17/2021).

[10] Hilje M Doekes, Glenn A Mulder, and Rutger Hermsen. “Repeated outbreaks drive the evolution of bacteriophage communication”. In: eLife 10 (Jan. 2021). Ed. by Samuel L Díaz-Muñoz, Aleksandra M Walczak, and Samuel L Díaz-Muñoz, e58410. issn: 2050-084X. doi: 10.7554/eLife.58410. url: https://doi.org/10.7554/eLife.58410 (visited on 04/30/2021).

[11] Zohar Erez, Ida Steinberger-Levy, Maya Shamir, Shany Doron, Avigail Stokar-Avihail, Yoav Peleg, Sarah Melamed, Azita Leavitt, Alon Savidor, Shira Albeck, Gil Amitai, and Rotem Sorek. “Communication between viruses guides lysis–lysogeny decisions”. In: Nature 541.7638 (Jan. 2017), pp. 488–493. issn: 1476-4687. doi: 10.1038/nature21049. url: https://doi.org/10.1038/nature21049.

[12] Ido Golding, Seth Coleman, Thu VP Nguyen, and Tianyou Yao. Decision Making by Temperate Phages. 2019. url: https://bacteriophysics.web.illinois.edu/wp/wp-content/uploads/2019/10/golding-encyclvirol-2019.pdf.

[13] Ellie Harrison and Michael A. Brockhurst. “Ecological and Evolutionary Benefits of Temperate Phage: What Does or Doesn’t Kill You Makes You Stronger”. eng. In: BioEssays: News and Reviews in Molecular, Cellular and Developmental Biology 39.12 (Dec. 2017). issn: 1521-1878. doi: 10.1002/bies.201700112.

[14] Martha M. Howe. “Bacteriophage Mu”. In: Molecular Microbiology. Ed. by Stephen J. W. Busby, Christopher M. Thomas, and Nigel L. Brown. Berlin, Heidelberg: Springer Berlin Heidelberg, 1998, pp. 65–80. isbn: 978-3-642-72071-0.

[15] Philippe Kourilsky. “Lysogenization by bacteriophage lambda”. In: Molecular and General Genetics MGG 122.2 (June 1973), pp. 183–195. issn: 1432-1874. doi: 10.1007/BF00435190. url: https://doi.org/10.1007/BF00435190.

[16] Guanlin Li, Michael H Cortez, Jonathan Dushoff, and Joshua S Weitz. “When to be temperate: on the fitness benefits of lysis vs. lysogeny”. In: Virus Evolution 6.2 (May 2020). veaa042. issn: 2057-1577. doi: 10.1093/ve/veaa042. eprint: https://academic.oup.com/ve/article-pdf/6/2/veaa042/33780375/veaa042.pdf. xurl: https://doi.org/10.1093/ve/veaa042.

[17] A. Lwoff. “Lysogeny”. In: Bacteriological reviews 17.4 (Dec. 1953). PMC180777[pmcid], pp. 269–337. issn: 0005-3678. url: https://pubmed.ncbi.nlm.nih.gov/13105613.

[18] Sergei Maslov and Kim Sneppen. “Well-temperate phage: optimal bet-hedging against local environmental collapses”. In: Scientific Reports 5.1 (June 2015), p. 10523. issn: 2045-2322. doi: 10.1038/srep10523. url: https://doi.org/10.1038/srep10523.

[19] Ron Milo. “What is the total number of protein molecules per cell volume? A call to rethink some published values”. In: Bioessays 35.12 (Dec. 2013), pp. 1050–1055. issn: 0265-9247. doi: 10.1002/bies.201300066. url: https://www.ncbi.nlm.nih.gov/pmc/articles/PMC3910158/ (visited on 05/11/2021).

[20] Amos B. Oppenheim, Oren Kobiler, Joel Stavans, Donald L. Court, and Sankar Adhya. “Switches in Bacteriophage Lambda Development”. en. In: Annual Review of Genetics 39.1 (Dec. 2005), pp. 409–429. issn: 0066-4197, 1545-2948. doi: 10.1146/annurev.genet.39.073003.113656. url: http://www.annualreviews.org/doi/10.1146/annurev.genet.39.073003.113656 (visited on 03/17/2021).

[21] Mark Ptashne. A Genetic Switch: Phage Lambda Revisited. en. CSHL Press, 2004. isbn: 9780879697167.

[22] Justin Silpe and Bonnie Bassler. “A Host-Produced Quorum-Sensing Autoinducer Controls a Phage Lysis-Lysogeny Decision”. In: Cell 176 (Dec. 2018). doi: 10.1016/j.cell.2018.10.059.

[23] F. M. Stewart and B. R. Levin. “The population biology of bacterial viruses: why be temperate”. In: Theor Popul Biol 26.1 (Aug. 1984), pp. 93–117.

[24] Lindi Wahl, Matthew Betti, David Dick, Tyler Pattenden, and Aaryn Puccini. “Evolutionary stability of the lysis-lysogeny decision: Why be virulent?” In: Evolution 73 (Nov. 2018). doi: 10.1111/evo.13648.

[25] Joshua S Weitz, Guanlin Li, Hayriye Gulbudak, Michael H Cortez, and Rachel J Whitaker. “Viral invasion fitness across a continuum from lysis to latency†”. In: Virus Evolution 5.1 (Apr. 2019). vez006. issn: 2057-1577. doi: 10.1093/ve/vez006. eprint: https://academic.oup.com/ve/article-pdf/5/1/vez006/30989561/vez006.pdf. url: https://doi.org/10.1093/ve/vez006.

[26] Joshua S. Weitz. Quantitative Viral Ecology: Dynamics of Viruses and Their Microbial Hosts. Princeton University Press, 2015. url: https://www.jstor.org/stable/j.ctt1dr36hx (visited on 04/17/2021).

[27] R. Young. “Bacteriophage lysis: mechanism and regulation”. In: Microbiological reviews 56.3 (Sept. 1992). PMC372879[pmcid], pp. 430–481. issn: 0146-0749. url: https://pubmed.ncbi.nlm.nih.gov/1406491.

[28] Lanying Zeng, Samuel O Skinner, Chenghang Zong, Jean Sippy, Michael Feiss, and Ido Golding. “Decision making at a subcellular level determines the outcome of bacteriophage infection”. In: Cell 141.4 (2010), pp. 682–691.

